# *Chlamydia trachomatis* suppresses host cell store-operated Ca^2+^ entry and inhibits NFAT/calcineurin signaling

**DOI:** 10.1101/2022.03.15.484523

**Authors:** Nicholas B. Chamberlain, Ted Hackstadt

## Abstract

The obligate intracellular bacterium, *Chlamydia trachomatis*, replicates within a parasitophorous vacuole termed an inclusion. During development, host proteins critical for regulating intracellular calcium (Ca^2+^) homeostasis interact with the inclusion membrane. The inclusion membrane protein, MrcA, interacts with the inositol-trisphosphate receptor (IP3R), an ER cationic channel that conducts Ca^2+^. Stromal interaction molecule 1 (STIM1), an ER transmembrane protein important for regulating store-operated Ca^2+^ entry (SOCE), localizes to the inclusion membrane via an uncharacterized interaction. We therefore examined Ca^2+^ mobilization in *C. trachomatis* infected cells. Utilizing a variety of Ca^2+^ indicators to assess changes in cytosolic Ca^2+^ concentration, we demonstrate that *C. trachomatis* impairs host cell SOCE. Ca^2+^ regulates many cellular signaling pathways. We find that the SOCE-dependent NFAT/calcineurin signaling pathway is impaired in *C. trachomatis* L2 infected HeLa cells and likely has major implications on host cell physiology as it relates to *C. trachomatis* pathogenesis.

## Introduction

The phylum *Chlamydiae* contains the human pathogens *Chlamydia trachomatis, C. psittaci*, and *C. pneumoniae*. Chlamydiae are obligate intracellular, gram negative bacteria that undergo a biphasic developmental cycle with the bacteria in either an infectious and metabolically constrained state, termed the elementary body (EB), or a metabolically active and replicative state that is non-invasive, classified as a reticulate body (RB) [1, 2]. The intracellular development of chlamydiae occurs within a parasitophorous vacuole that is referred to as an inclusion [3]. *C. trachomatis*, contains multiple serological variants that demonstrate specific organotropism and associated disease. Serovars A-C are associated with endemic, blinding trachoma, serovars D-K are responsible for the common sexually transmitted urogenital infections, and serovars L1-L3 are the causative agents of lymphogranuloma venereum, a more invasive disease [4-6].

*C. trachomatis* utilizes a type III secretion system to deliver effectors across the inclusion membrane and into the cytosol of the host cell [7]. A subset of these T3SS effector proteins, termed Incs, localize to the inclusion membrane via a bi-lobed transmembrane domain [8, 9]. Incs are oriented in the inclusion membrane in a manner that exposes them to the cytosol of the host cell [10, 11], enabling the Incs to interact with cytosol-exposed proteins [12]. While Incs are typically distributed relatively evenly around the inclusion membrane, a subset of Incs are localized to discrete, punctate sites on the membrane, termed microdomains, that are also enriched in cholesterol and active host Src-family kinases [13]. Currently, the *C. trachomatis* effectors CT101 (MrcA), CT147, CT222, CT223 (IPAM), CT224, CT228, CT232 (IncB), CT233 (IncC), CT288, and CT850 have been identified as Incs enriched in inclusion microdomains [9, 13-15]. Early findings demonstrated microdomains function as hubs for interactions with the cytoskeleton and promote the positioning of the inclusion at the microtubule organizing center [13, 16]. Microdomains also influence extrusion-based dissemination, a process in which all or part of an intact chlamydial inclusion is exocytosed from the host cell for dissemination to distal anatomic sites. The phosphorylation of myosin light chain 2 (MLC2) at the microdomain is a critical regulator of this dissemination process [14], which occurs via Ca^2+^-dependent pathways [17].

Cytosolic Ca^2+^ acts as a second messenger for a number of pathways in eukaryotes influencing fertilization, embryonic axis formation, cell differentiation, cell proliferation, transcription factor activation, and cell fate decisions [18, 19]. Cytosolic Ca^2+^ is tightly regulated to ensure these pathways are activated only when required. This homeostatic regulation of cytosolic Ca^2+^ concentration is achieved through Ca^2+^ pumps, exchangers, channels, buffering proteins, and sensors to control Ca^2+^ movement across the plasma membrane and the membranes of intracellular Ca^2+^ stores [20]. In excitable and non-excitable cells, store-operated Ca^2+^ entry (SOCE) is a major regulator of Ca^2+^ homeostasis [21]. The SOCE process is initiated by the depletion of ER Ca^2+^. The reduction in ER luminal Ca^2+^ is sensed by stromal interaction molecule 1 (STIM1) and subsequently initiates a STIM1 conformational change enabling STIM1 to interact with Orai1 to generate Ca^2+^ release-activated (CRAC) influx channels at the plasma membrane. The STIM1-CRAC channel interaction opens the channel resulting in an ingress of extracellular Ca^2+^ into the cytosol [22]. This influx of extracellular Ca^2+^ into the cytosol refills Ca^2+^ stores and is a signal for select Ca^2+^-dependent pathways.

The *C. trachomatis* inclusion interacts with the host ER, a major intracellular Ca^2+^ store, through multiple interactions including the inclusion membrane protein IncD interaction with ceramide transfer protein (CERT) and the binding of CERT to VAMP-associated proteins (VAP), ER resident proteins [23], the direct interaction of IncV with VAP [24], the interaction of STIM1 to the inclusion membrane by an unknown mechanism [17, 25], and the interaction of the inclusion membrane protein MrcA with the inositol triphosphate receptor (IP_3_R) [17]. STIM1 and MrcA localization to microdomains of the inclusion membrane [17, 25], demonstrate that regulators of Ca^2+^ homeostasis are actively subverted during chlamydial development. Therefore, we investigated if chlamydial development disrupts host cell Ca^2+^ homeostasis. Here we provide evidence that SOCE is impaired by the midpoint of the chlamydial developmental cycle and the SOCE-dependent NFAT/calcineurin signaling pathway is concurrently abrogated.

## Results

### Store-operated Ca^2+^ entry is impaired by mid-cycle of the *C. trachomatis* developmental cycle

To investigate the impact of chlamydial infections upon intracellular Ca^2+^ mobilization, the ratiometric Ca^2+^ indicator, Fura-2, AM, was used to assess changes in host cell intracellular Ca^2+^ concentration ([Ca^2+^]_i_). For this analysis of intracellular Ca^2+^ mobilization, HeLa cells either uninfected or infected with *C. trachomatis* L2 were loaded with Fura-2, AM at the desired time post infection. The binding of Ca^2+^ to Fura-2, AM induces a shift in its fluorescence excitation from 380 nm to 340 nm. Therefore, a 340nm/380nm fluorescence ratio of Fura-2 was used to obtain relative changes in cytoplasmic [Ca^2+^]_i_. A Ca^2+^ re-addition assay [26] was performed to quantify relative changes in [Ca^2+^]_i_ caused by Ca^2+^ mobilization events. Ratiometric measurements were taken when cells were in a resting state, during induced ER Ca^2+^ leakage, and during SOCE. Following a resting state baseline reading in Ca^2+^-free Ringer’s solution, cells were incubated with Ca^2+^-free Ringer’s solution containing either thapsigargin (TG) or the vehicle control, DMSO. TG is an inhibitor of sarco-endoplasmic reticulum Ca^2+^ ATPase (SERCA), a Ca^2+^ pump responsible for mobilizing Ca^2+^ from the cytosol into the lumen of the ER or sarcoplasmic reticulum. TG thus causes an increase in cytosolic Ca^2+^ by impairing ER Ca^2+^ uptake while Ca^2+^ egress from the ER continues to occur via passive ER Ca^2+^ leakage through ER translocon complexes [27, 28].

The calcium re-addition assay was performed to assess changes in [Ca^2+^]_i_ for uninfected and infected HeLa cells. When TG treated and DMSO control HeLa cells were moved to a Ca^2+^-containing Ringer’s solution, a distinctive increase in [Ca^2+^]_i_ was detected in TG-treated cells, but not DMSO treated, indicating that Ca^2+^ depletion of the ER resulted in the activation of SOCE (Fig 1A). A STIM1 siRNA knockdown was performed to verify that this methodology was able to detect impaired SOCE (Extended Data Fig. 1).

**Fig 1:**
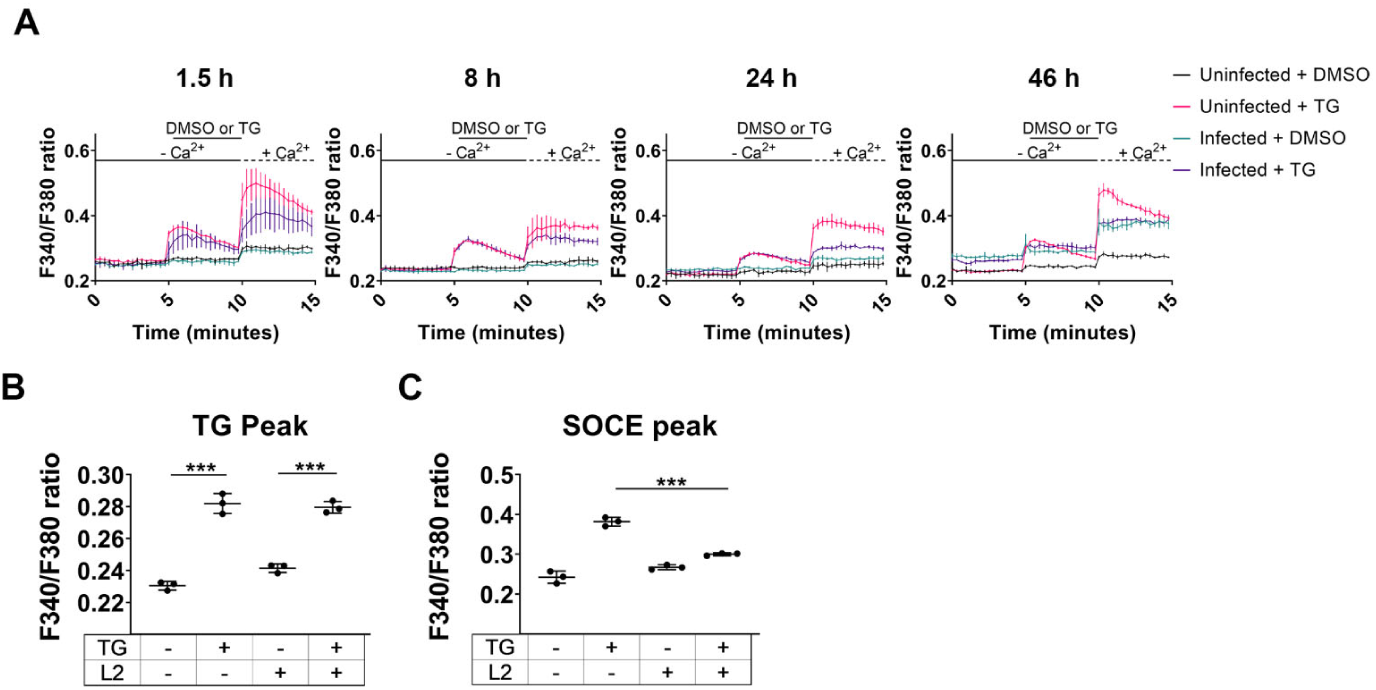
*C. trachomatis* impairs host cell store-operated calcium entry (SOCE) by a mid-cycle developmental time point. a Ca^2+^ re-addition assays were performed with Fura-2, AM to assess [Ca^2+^]_i_ changes in HeLa cells infected with *C. trachomatis* L2 or uninfected. Thapsigargin (TG) was used to induce ER Ca^2+^ depletion and subsequent SOCE. DMSO was used as a vehicle control. A ratiometric assessment of Fura-2, AM fluorescence at 340 nm and 380 nm was taken to calculate the relative change in [Ca^2+^]_i_. b The peak calcium efflux from the ER induced by TG for each condition at 24 hpi was measured. The TG treatment was compared to DMSO for either uninfected or infected cells. Student’s T-test was used to compare the DMSO to the TG treatment, n=3. c Peak SOCE for each sample at the 24 hpi timepoint. Student’s T-test was used to compare the uninfected TG treated condition to the 24 hpi TG treated condition, n=3. Data (**a-c**) are presented as mean ± SEM.

To interrogate changes in Ca^2+^ mobilization during *C. trachomatis* development, the Fura-2, AM Ca^2+^ re-addition assay was performed with HeLa cells infected with *C. trachomatis* L2 at early-(1.5 hpi), early-to-mid-(8 hpi), mid (24 hpi)-, and late-(46 hpi) cycle developmental time points. Similar TG-induced Ca^2+^ egress from the ER was observed in infected cells vs uninfected at the mid-cycle timepoint (Fig 1b). However, the TG-induced SOCE change in [Ca^2+^]_i_ was severely impaired in infected cells at a mid-cycle developmental time point compared to the uninfected (Fig 1c). At 46 hpi, the baseline level of F340/F380 was elevated in the infected vs the uninfected. The likely interpretation of this would be that membranes were compromised due to lysis resulting in elevated [Ca^2+^]_i_ at this timepoint, which may indicate why TG-induced ER Ca^2+^ egress and SOCE at this time point were impaired (Fig 1a).

### Verification of *C. trachomatis*-suppressed SOCE using single-cell analysis

The Fura-2, AM-based method measured changes in cytoplasmic Ca^2+^ at the cell population level. To assess the influence of *C. trachomatis* serovar L2 on host cell Ca^2+^ mobilization at a single cell level, we used the Ca^2+^ indicator Fluo-4, AM with live-cell microscopy to determine changes in [Ca^2+^]_i_ in infected and uninfected cells. To quantify the normalized relative change in Fluo-4, AM fluorescence intensity (F) for each cell, ΔF/F_0_ was calculated. The baseline resting state fluorescence (F_0_) was the average of the first four mean intensity measurements for the ROI in Ca^2+^-free Ringer’s solution, and ΔF = F-F_0_ was used to calculate the change in fluorescence.

Analysis of [Ca^2+^]_i_ mobilization in uninfected and *C. trachomatis*-infected cells was performed at a mid-cycle (24 hr) developmental timepoint. In uninfected and infected HeLa cells treated with the DMSO carrier, stochastic Ca^2+^ elevations were observed in a small subset of cells, and when DMSO-treated control and infected cells were placed in Ca^2+^-Ringer’s solution, there was a modest increase in [Ca^2+^]_i_. When cells were treated with TG, a dramatic increase in ΔF/F_0_ was observed for both uninfected and infected cells, indicating the TG treatment mobilized Ca^2+^ from the ER into the cytoplasm. The transition of TG-treated cells to Ca^2+^-containing Ringer’s solution resulted in a striking increase in ΔF/F_0_ associated with SOCE in uninfected but not infected cells (Fig 2a). The mean of the single-cell measurements provides a visualization of the single cell Ca^2+^ flux trends for each condition (Fig 2b). TG-treated, *C. trachomatis*-infected cells demonstrated a significant increase in relative fluorescence of the Fluo-4, AM indicator compared to uninfected TG-treated cells (Fig 2c). At 44 hpi, there was no significant difference in the TG peak between TG-treated uninfected and infected cells. (Extended Data Fig. 2a-c). These results indicate that TG induces Ca^2+^ egress from the ER of uninfected and infected cells.

**Fig 2:**
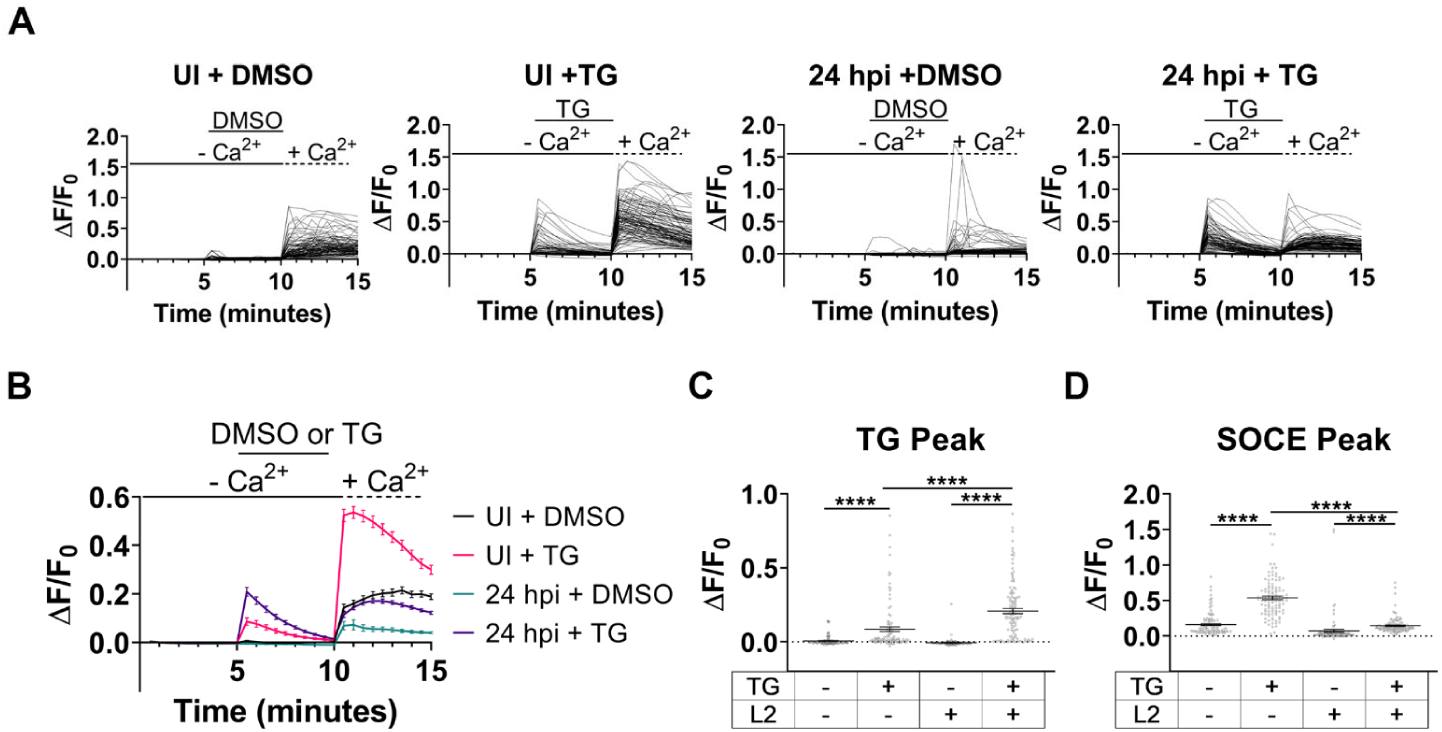
Fluo-4 assessment of *C. trachomatis* suppression of host cell SOCE. Fluo-4, AM, was used with live-cell microscopy to assess cytosolic calcium mobilization in uninfected and *C. trachomatis* L2-infected cells. a Single-cell analysis of the relative change in Fluo-4 fluorescence during Ca^2+^ re-addition assay was performed for uninfected and *C. trachomatis*-infected cells at a mid-cycle developmental timepoint. For each condition, ≥ 39 cells were analyzed. b The mean relative change in Fluo-4 fluorescence was calculated for each condition from A. c The relative change in Fluo-4 fluorescence was assessed at the peak TG-induced ER Ca^2+^ egress. d The relative change in Fluo-4 fluorescence was calculated for peak SOCE. A Kruskal-Wallis test was performed with Dunn’s post-hoc multiple comparisons test to compare the conditions (**c-d**). Comparisons denoted with **** have a p value <0.0001 and ns represents no significant difference. Data (**b-d**) are presented as mean ± SEM.

The SOCE of uninfected and infected cells was also assessed using the Fluo-4, AM indicator. The mean ΔF/F_0_ at the SOCE peak for HeLa cells infected with *C. trachomatis* serovar L2 was severely reduced compared to uninfected cells. Furthermore, the uninfected, TG-treated cells had 53% of the cells with a ΔF/F_0_ greater than 0.5 while infected cells only had 3% of the cells with a ΔF/F_0_ greater than 0.5 (Fig 2d). Additionally, uninfected and infected DMSO treated cells had 5% and 2% of cells, respectively, with a ΔF/F_0_ of greater than 0.5 (Fig 2d). At 44 hpi, infected cells had a significantly reduced ΔF/F_0_ mean at the SOCE peak compared to uninfected, and only 3% of the cells had a SOCE peak greater than 0.5 in the infected and TG-induced condition compared to 37% in the uninfected and TG-induced condition (Extended Data Fig. 2d). Collectively, the Fluo-4, AM single-cell analysis demonstrated that SOCE of the host cell is impaired by mid-cycle and remains suppressed at later developmental time points.

### Genetically encoded Ca^2+^ indicator confirms SOCE inhibition in *C. trachomatis* infected cells

The genetically encoded Ca^2+^ indicator (GECI), GCaMP6m, was used to corroborate the results of the Fura-2 AM and Fluo-4, AM based quantification of [Ca^2+^]_i_ mobilization in uninfected and *C. trachomatis*-infected cells. The GCaMP family of GECIs are single fluorescent protein indicators that operate by utilizing a circularly permuted GFP linked to calmodulin and the Ca^2+^-dependent calmodulin-interacting peptide M13 from myosin light chain kinase (MLCK). Ca^2+^ binding induces a conformational change in the circularly permuted GFP causing a large increase in fluorescence intensity [29]. Utilizing GCaMP6m, a relative change in GFP fluorescence can be performed to determine changes in [Ca^2+^]_i_. *C. trachomatis* L2 expressing mScarlet, permitted visualization of chlamydial inclusions during imaging. HeLa cells infected with mScarlet *C. trachomatis* serovar L2 and transfected with pN1-GCaMP6m-XC were tested to assess Ca^2+^ mobilization.

The single-cell and mean analysis of the relative fluorescence change of GCaMP6m demonstrated similar trends as the Fluo-4 analysis with SOCE of infected cells impaired relative to uninfected cells (Fig 3a,b). The single-cell analysis at the TG peak indicated that both uninfected and infected cells had elevated fluorescence upon TG addition compared to the DMSO vehicle control (Fig 3c). The single-cell analysis of the SOCE peak indicated that uninfected cells treated with TG had an increased relative change in GCaMP6m fluorescence compared to the DMSO control, however, there was no significant difference in the SOCE peak between the 24 hpi cells treated with DMSO and TG. (Fig 3d). The GCaMP6m analysis of HeLa cells at a mid-to-late developmental cycle timepoint, 36 hpi, with *C. trachomatis* serovar L2 demonstrated that SOCE was suppressed in infected cells (Extended Data Fig. 3). The Fura-2, Fluo-4, and GCaMP6m Ca^2+^ indicators collectively demonstrated that SOCE is impaired in HeLa cells infected with *C. trachomatis* L2 by mid-cycle developmental time point.

**Fig 3:**
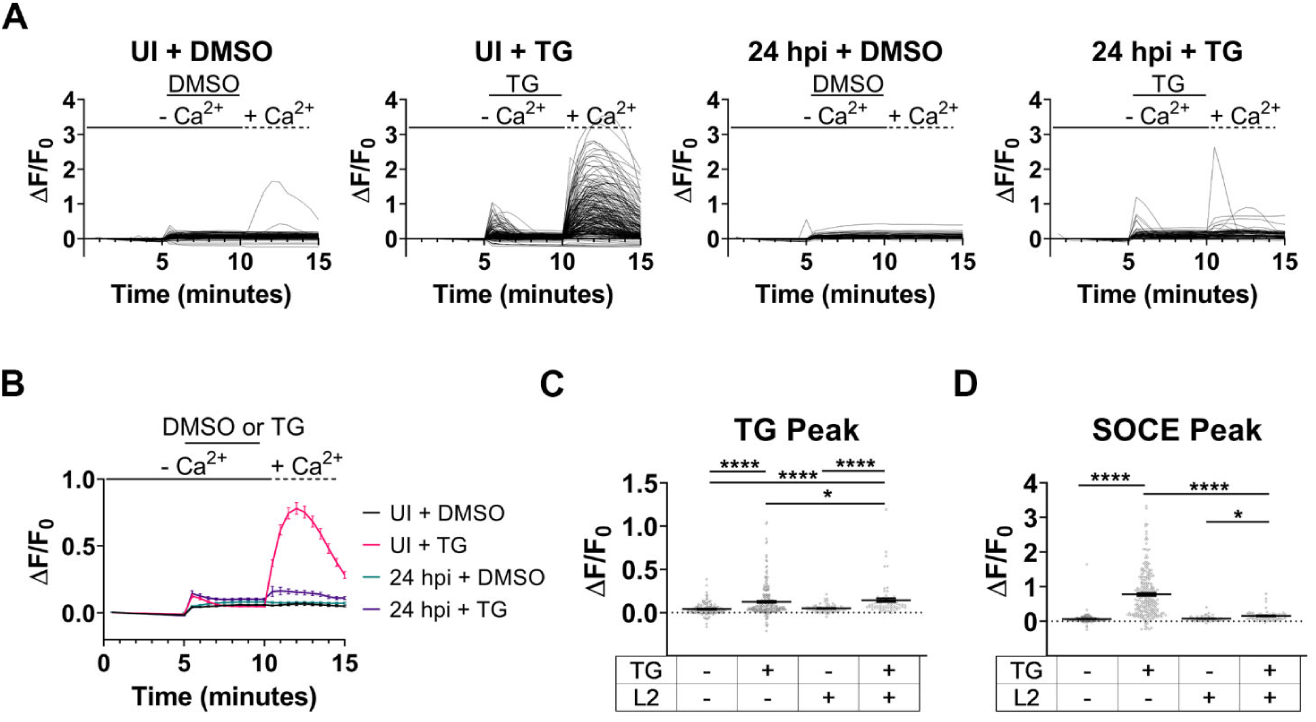
Genetically encoded Ca^2+^ indicator verification of impaired SOCE of host cell. Live-cell microscopy with the genetically encoded Ca^2+^ indicator, GCaMP6m, was used to measure changes in [Ca^2+^]_i_. The Ca^2+^ re-addition assay was performed with *C. trachomatis-*infected and uninfected HeLa cells. a Relative change in GCaMP6m fluorescence was calculated throughout Ca^2+^ re-addition assay for individual HeLa cells at a mid-cycle (24 h) developmental timepoint.. For each condition, ≥ 67 cells were analyzed. b The mean relative change of GCaMP6m fluorescence was measured from the single-cell analysis in a. c The relative change in GCaMP6m fluorescence at peak TG-induced ER Ca^2+^ efflux was calculated. d The relative change in GCaMP6m fluorescence was measured at peak ΔF/F_0_ during SOCE. Because GCaMP6m measurements were a non-Gaussian data set, a Kruskal-Wallis test was performed with Dunn’s post-hoc multiple comparisons test to compare the conditions (**c**,**d**). Comparisons denoted with **** have a p value <0.0001 and * have a p value <0.05. Data (**b-d**) are presented as mean ± SEM.

To determine if a urogenital *C. trachomatis* serovar also impairs SOCE of the host cell, the Ca^2+^ re-addition assay was performed using GCaMP6m in HeLa cells infected with *C. trachomatis* serovar D at the mid-cycle (24 hr) developmental timepoint. Similar to the *C. trachomatis* L2 results, *C. trachomatis* D impaired SOCE of the host cell by the mid-cycle developmental timepoint (Extended Data Fig. 4).

### *C. trachomatis* prevents SOCE-induced NFAT nuclear localization

Extended increases in [Ca^2+^]_i_ caused by SOCE result in the activation of various Ca^2+^-dependent pathways. To gain insights into physiological consequences of impaired SOCE in C. trachomatis infected cells, a specific SOCE-dependent pathway was investigated. Sustained elevated [Ca^2+^]_i_ activates the calcineurin-NFAT signaling pathway via the Ca^2+^-mediated binding of calmodulin to calcineurin, the calcineurin-dependent dephosphorylation of the NFAT transcription factor to expose its nuclear localization signal, and the subsequent cytosol-to-nucleus translocation of NFAT. We investigated the SOCE-induced nuclear localization of NFAT1 in *C. trachomatis*-infected cells at the mid-cycle developmental time point to determine if a SOCE-dependent pathway is abrogated during *C. trachomatis* infection. HeLa cells infected with mScarlet-expressing *C. trachomatis* L2 were transfected with the HA-NFAT1(4-460)-GFP plasmid. A pilot study indicated that the optimal time following TG treatment and Ca^2+^ incubation for NFAT-GFP nuclear translocation in HeLa cells was 18 min post-Ca^2+^ addition (Supplementary Video 1). Therefore, imaging for NFAT-GFP nuclear translocation was performed immediately following TG or DMSO treatment, and again at 18 min post-Ca^2+^ incubation. Imaging of NFAT-GFP-expressing HeLa cells treated with DMSO demonstrated no noticeable change in nuclear NFAT-GFP fluorescence following the 18 min Ca^2+^ incubation, while the TG treatment caused a dramatic increase in nuclear NFAT-GFP following the Ca^2+^ incubation (Fig 4a). However, neither the DMSO nor TG treatment caused a noticeable increase of nuclear NFAT-GFP in *C. trachomatis*-infected cells at 24 hpi (Fig 4a). A representative time course video of NFAT-GFP-expressing HeLa cells infected with mScarlet *C. trachomatis* L2 did not demonstrate NFAT-GFP nuclear localization in the presence of TG, however, NFAT-GFP-expressing cells in the same field of view without a chlamydial inclusion showed nuclear translocation of NFAT-GFP (Supplementary Video 2).

**Fig 4:**
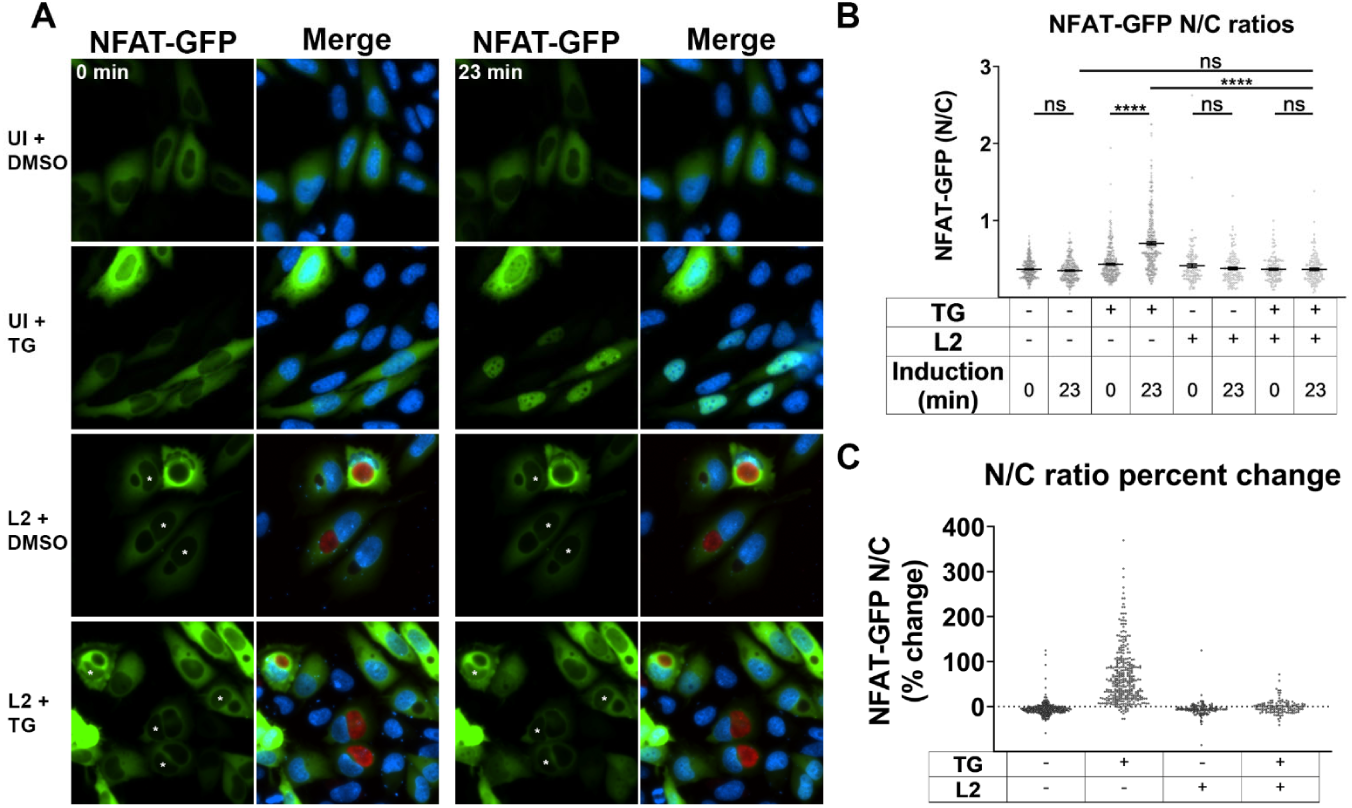
NFAT nuclear localization is diminished in *C. trachomatis*-infected cells. Nuclear translocation of NFAT-GFP was assessed in *C. trachomatis* L2 infected and uninfected HeLa cells. a Live-cell imaging was performed with HeLa cells transfected with NFAT-GFP and either infected with *C. trachomatis* or uninfected. Cells were induced with thapsigargin (TG) to trigger SOCE or treated with the DMSO vehicle control for 5 min. Cells were imaged at 0 min post-addition of TG or DMSO, and then imaged at 18 min after administering Ca^2+^-containing Ringer’s solution. Asterisks denote the nuclei of *C. trachomatis*-infected cells. b Single-cell analysis of the NFAT-GFP nuclear to cytoplasmic mean fluorescence ratio (N/C) was calculated for each cell immediately following TG or DMSO treatment, and the NFAT-GFP N/C ratio was calculated again following the 18 min Ca^2+^ incubation. For each condition, ≥ 134 cells were analyzed. A Kruskal-Wallis test was performed with Dunn’s post-hoc multiple comparisons test. Comparisons denoted with **** have a p value <0.0001 and ns represents no significant difference. The data are presented as mean ± SEM. c The percent change in NFAT-GFP N/C ratio was calculated for the N/C ratios from (**b**) for each individual cell.

Image analysis was performed to calculate the NFAT-GFP nuclear to cytoplasmic fluorescence ratio (N/C) for each condition. The NFAT-GFP N/C was measured as the mean fluorescence intensity of nuclear NFAT-GFP / the mean fluorescence intensity of cytoplasmic NFAT-GFP for individual cells. Cells with an observable inclusion containing mScarlet *C. trachomatis* were calculated for infected cells. The only significant difference identified between the pre- and post-incubation with Ca^2+^-containing Ringer’s solution was for the uninfected + TG condition, which demonstrated a substantial increase in the NFAT-GFP N/C ratio following the 18 min Ca^2+^ solution incubation (p value < 0.0001). Thus, no significant difference was seen for *C. trachomatis*-infected cells between post-TG treatment and post-Ca^2+^ treatment. Additionally, no difference was observed between the NFAT-GFP N/C ratios of the uninfected + DMSO condition post-Ca^2+^ treatment and the 24 hpi *C. trachomatis* infection + TG condition post-Ca^2+^ incubation. The NFAT-GFP N/C ratio of the 24 hpi *C. trachomatis* + TG condition post-Ca^2+^ incubation was drastically reduced compared to the uninfected + TG condition post-Ca^2+^ incubation (Fig 4b). To visualize how the NFAT-GFP N/C ratio changed from post-TG treatment to post-Ca^2+^ incubation for each cell, a percent change in NFAT-GFP N/C ratio was calculated per cell (Fig 4c). Collectively, the NFAT-GFP N/C ratio assessment demonstrated a severe reduction in NFAT-GFP nuclear localization in *C. trachomatis* serovar L2 infected HeLa cells when induced to undergo SOCE.

## Discussion

We examined Ca^2+^ dynamics in *C. trachomatis*-infected cells and demonstrate that *C. trachomatis* inhibits SOCE of the host cell by a mid-cycle (24 h) developmental time point. This inhibition of SOCE was confirmed using three independent methods of intracellular Ca^2+^ quantitation. High concentrations of intracellular Ca^2+^ resulting from SOCE induction can act as a messenger to activate numerous signaling pathways [18, 30]. Among the pathways activated is the calcineurin/NFAT pathway. Sustained high concentrations of intracellular Ca^2+^ activates the phosphatase calcineurin which, in turn, dephosphorylates the cytoplasmic components of the NFAT transcription complex to trigger NFAT translocation into the nucleus [31]. NFAT transcription complexes regulate genes encoding immunomodulatory proteins or involved in developmental cellular differentiation [19, 32, 33]. *C. trachomatis* inhibition of SOCE had the downstream effect of inhibiting the calcineurin/NFAT pathway, thus providing a biological confirmation of the intracellular Ca^2+^ quantitation. The impairment of NFAT1 nuclear translocation demonstrate a specific signaling pathway that is affected by suppressing SOCE in *C. trachomatis* infected cells and likely has a multifaceted impact on host cell physiology and chlamydial pathogenesis.

At the end of their developmental cycle, chlamydiae are released by one of two distinct mechanisms for infection of adjacent cells and subsequent cycles of infection. Infectious EBs are released either by lysis of the infected cell, or intact or partially intact membrane bound inclusions are released by a process known as extrusion [34]. The cellular requirements and mechanisms for the two different release mechanisms are unique. Myosin II and Ca^2+^ are essential for chlamydial extrusion-based dissemination [14, 17, 34]. Ca^2+^ appears to be an important determinant of mechanism for chlamydial release from the host cell, however, the suppression of SOCE indicates that other sources of Ca^2+^ must be utilized.

Although SOCE does not appear to be the source of Ca^2+^ required for extrusion, host proteins involved in Ca^2+^ homeostasis are recruited to inclusion membrane microdomains that serve as a hub for interactions with the cytoskeleton and the ER [13, 35, 36]. The microdomain inclusion membrane protein, MrcA, interacts with the Ca^2+^ channel, IP_3_R [17]. The activation of IP_3_R depletes ER Ca^2+^ and typically results in STIM1 formation of plasma membrane-localized CRAC channels and SOCE induction [21]. STIM1 is a resident ER Ca^2+^ sensor that detects ER Ca^2+^ depletion and acts, in conjunction with Orai1, to trigger SOCE. The knockdown of STIM1 dampens the amount of Ca^2+^ mobilized through IP_3_R [37] and impairs extrusion production [17], thus it appears that *C. trachomatis* mobilizes Ca^2+^ from the ER to provide the Ca^2+^ necessary for activating extrusion machinery. Ca^2+^ can exhibit steep concentration gradients even within a single cell [30]. An elevated Ca^2+^ microenvironment spatially resolved to the inclusion microdomain-ER interface might help explain the asynchrony and lack of uniformity to the extrusion process. The lack of synchrony and the fact that all infected cells do not undergo extrusion may present challenges in characterizing the formation of Ca^2+^ microenvironments near the inclusion.

Based upon these observations, we propose a working model in which IP_3_R activation is important for elevating cytosolic Ca^2+^ near the inclusion microdomains, and STIM1 ensures optimal efflux of Ca^2+^ through IP_3_R. Furthermore, we suggest that STIM1 interaction with the inclusion prevents its diffusion in the ER during Ca^2+^ depletion, and thus precludes STIM1 localization to ER-PM junctions and the activation of CRAC channels essential to SOCE (Fig 5).

**Fig 5:**
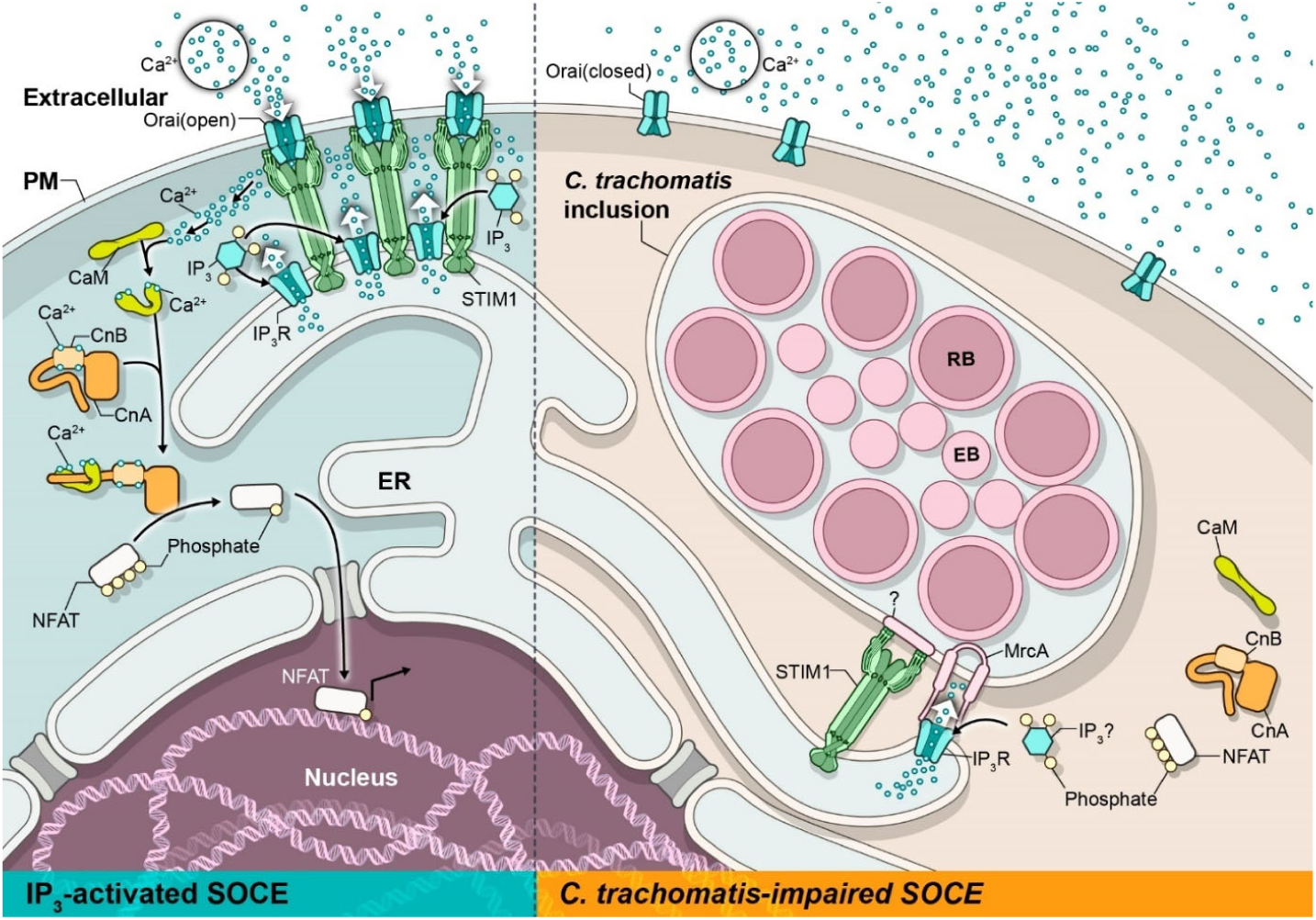
Working model for *C. trachomatis* impaired SOCE. During inositol-trisphosphate (IP_3_)-induced SOCE (left side), IP_3_ binds to IP_3_R and triggers the egress of Ca^2+^ from the ER to the cytoplasm through IP_3_R. The depletion in ER luminal Ca^2+^ is recognized by the EF-hand domain of STIM1, resulting in a conformational change of STIM1. This enables the STIM1 CAD domain to interact with Orai1 at the PM to form Ca^2+^ release-activated Ca^2+^ (CRAC) channels. The opening of CRAC channels results in an influx of extracellular Ca^2+^ into the cytosol of the cell, which refills the ER Ca^2+^ store as well as acts as a signal for Ca^2+^-dependent proteins. The Ca^2+^-dependent protein, calcineurin, is activated by SOCE. Calcineurin dephosphorylation of NFAT exposes the nuclear localization signal, resulting in NFAT nuclear import to regulate expression of various genes. Our current working model for *C. trachomatis* impaired SOCE (right side) hypothesizes that the sequestering of the STIM1 CAD domain at the inclusion membrane microdomain prevents STIM1 from interacting with Orai1 to induce SOCE. We hypothesize that the MrcA-IP_3_R interaction [17] leads to the activation of IP_3_R and mobilization of ER Ca^2+^ into the cytosol near the inclusion microdomain. Since SOCE is impaired in *C. trachomatis*-infected cells, efflux of ER Ca^2+^ would not elicit SOCE. *C. trachomatis* suppression of SOCE prevents the SOCE-dependent activation of calcineurin/NFAT signaling of the host cell and likely other SOCE-dependent pathways.

The primary sites of *C. trachomatis* infection for genital infections in women and ocular infections are the endocervical epithelium and conjunctival epithelium, respectively [4]. As the first targets of infection, mucosal epithelial cells act as first responders to pathogen challenge by the secretion of cytokines and chemokines, and thus play a key role in the innate immune response [38]. In an immortalized primary human endocervix-derived epithelial cell line, a productive *C. trachomatis* infection mitigated a pro-inflammatory cytokine and chemokine response. Although the mechanism was not defined, it was proposed that the circumvention of a robust cytokine and chemokine response represented a potential evasion strategy promoting the establishment of a favorable intracellular niche within the endocervix epithelium [39]. SOCE impacts cytokine and chemokine signaling events [30]. The influence of *C. trachomatis*-mediated suppression of SOCE warrants further investigation.

The impairment to calcineurin activation and NFAT1 nuclear translocation demonstrate a specific signaling pathway that is affected by chlamydial suppression of SOCE. Although the major functions of NFAT are often considered in innate immune cells such as lymphocytes, macrophages, dendritic cells, or neutrophils, NFAT plays multiple additional roles in development and cellular differentiation [19]. NFAT is also expressed in multiple cell types [19], including epithelial cells ([40-42] and endothelial cells [43], however, direct information on how disruption of NFAT signaling might impact pathogen interaction with epithelial cells is limited. African swine fever virus inhibits NFAT-regulated transcription of immunomodulatory proteins by synthesis of a protein, A238L, that directly binds to and inhibits the calcineurin phosphatase activity required for NFAT activation [44]. Reactivation of latent Epstein-Barr virus to a lytic state via a Ca^2+^/calcineurin dependent activation of NFAT by a complex mechanism has been proposed to represent a negative feedback loop involving a viral protein, Zta, that directly binds to and attenuates NFAT [45]. *Helicobacter pylori* is associated with peptic ulcers and gastric adenocarcinoma. *H. pylori* has the capacity to influence NFAT activity either positively or negatively by the bacterial proteins CagA or VacA, respectively, in gastric epithelial cells [42]. VacA forms an anion selective channel in the plasma membrane [46], which is believed to deregulate membrane depolarization and inhibit SOCE. Although the mechanism of SOCE inhibition differs, *C. trachomatis* also inhibits SOCE and NFAT nuclear translocation. We hypothesize that this inhibition of NFAT signaling promotes chlamydial survival, possibly by downregulation of chemokine or cytokine production. Further studies in human primary cervical epithelial cells or animal model systems will undoubtedly be necessary to fully elucidate the benefits to chlamydial and host survival.

## Materials and methods

### Bacterial and mammalian cell culture

HeLa 229 human cervical epithelial-like cells (American Type Culture Collection) were cultivated in RPMI-1640 (Gibco) supplemented with 5% fetal bovine serum (HyClone) at 37°C and 5% CO_2_ in a humidified incubator. The *C. trachomatis* D (UW-3-Cx) and L2 (LGV 434) serovars were cultured in HeLa cells and EBs purified by density gradient centrifugation as previously described [47]. Infectious titers were determined by Inclusion Forming Unit (IFU) assays [48] with inclusions detected by indirect immunofluorescence.

### Cell transfection

For siRNA transfections, HeLa cells were seeded at 1×10^4^ cells/well in 96-well, black-wall, clear-bottom plates (Costar). Transfection complexes, either ON-TARGETplus Non-Targeting Control Pool siRNA (Dharmacon) or ON-TARGETplus SMARTpool STIM1 siRNA (Dharmacon) were diluted to a 1 μM working concentration in Opti-MEM. An equivalent volume of the diluted DharmaFECT 1 reagent and diluted siRNA were combined and incubated at room temperature for 30 min to form siRNA/DharmaFECT complexes. For the mock transfection, DharmaFECT 1 reagent in Opti-MEM was used. HeLa growth medium was exchanged with Opti-MEM while siRNA transfection complexes were generated. Following incubation, 0.1 mL of the diluted transfection complex was added per well. Cells were incubated for 48 hours prior to cell infection.

For mammalian expression vector transfections, HeLa cells that were either infected with *C. trachomatis*, L2, for the stated times or uninfected were transfected with the desired mammalian expression vector using X-tremeGENE HP DNA transfection reagent (Sigma) according to the manufacturer’s instructions.

### Fura-2, AM Ca^2+^ measurement

At the desired time post infection, medium was removed and cells were washed with PBS. Cells were loaded with 5 μM Fura-2, AM in Ca^2+^-containing Ringer’s Solution (140 mM NaCl, 5 mM KCl, 1 mM MgCl_2_, 10 mM HEPES, 2 mM CaCl_2_, 10 mM glucose, pH 7.4) for 1 hour at room temperature. Cells were washed twice with PBS and incubated in Ca^2+^-free Ringer’s Solution (140 mM NaCl, 5 mM KCl, 1 mM MgCl_2_, 10 mM HEPES, 3 mM EGTA, 10 mM glucose, pH 7.4) for 30 min at room temperature. Following incubation, Fura-2, AM readings were performed by measuring fluorescence at 340/11 nm and 380/20 nm excitation and 508/20 emission every 20 seconds for 5 minutes using a Cytation 5 (BioTek). The solution was replaced with Ca^2+^-free Ringer’s solution containing either 2 μM thapsigargin (TG) or an equivalent volume of DMSO (vehicle control). Fluorescence readings were repeated every 20 sec for 5 min. Each solution was then exchanged with Ca^2+^-containing Ringer’s solution, and fluorescence readings were taken every 20 sec for 5 min. Each condition was performed in triplicate.

### Fluo-4, AM Ca^2+^ measurement

At the desired time post-infection, cultures were washed with PBS and loaded with 2.5 μM Fluo-4, AM in Ca^2+^-containing Ringer’s Solution for 1 hour at room temperature. Cells were washed twice with PBS and incubated in Ca^2+^-free Ringer’s solution (140 mM NaCl, 5 mM KCl, 3 mM MgCl_2_, 10 mM HEPES, 10 mM glucose, pH 7.4) for 30 min at room temperature. Following incubation, cells were imaged using the Nikon Ti2e with a CFI60 Super Plan Fluor Phase Contrast ADM ELWD 40x Objective Lens (N.A. 0.6,; Nikon). ND acquisition (NIS-Elements 64-bit version 5.11.02 software (Nikon) was set to acquire images from 4 locations per well for infected and uninfected condition. ND acquisition was programmed to image every 30 seconds for 5 minutes. At the end of the Ca^2+^-free buffer imaging, the Ca^2+^-free buffer was exchanged with Ca^2+^-free buffer containing either 2 μM thapsigargin (TG) or DMSO carrier. Cells were imaged for another 5 min imaging interval. At the end of that interval, buffers were exchanged with a Ca^2+^-containing Ringer’s solution (140 mM NaCl, 5 mM KCl, 1 mM MgCl_2_, 10 mM HEPES, 3 mM EGTA, 10 mM glucose, pH 7.4), and images aquired for another 5 min interval. Huygens Essential software version 20.04 (Scientific Volume Imaging) was used to stabilize frames between time points and Imaris x64 software version 9.6.0 (Oxford Instruments) was used to measure the mean fluorescence intensity of Fluo-4 per cell at each time point.

### GCaMP6m Ca^2+^ measurement

HeLa cells that were either uninfected or infected with mScarlet-expressing *C. trachomatis* L2 were transfected with pGP-CMV-GCaMP6m. The pGP-CMV-GCaMP6m plasmid was a gift from Douglas Kim and the GENIE project (Addgene plasmid # 40754; RRID:Addgene_40754) [49]. At the desired time post infection, medium was removed from cells, and cells were washed with PBS. Cells were incubated in Ca^2+^-free Ringer’s solution for 15 min. Imaging, solution exchanges, image processing, and fluorescence measurements were performed as described in the Fluo-4, AM Ca^2+^ measurements section. Imaging was performed at 37°C, 5% CO_2_, 92% relative humidity using a stage-top incubation chamber (Okolab).

### NFAT1-GFP nuclear translocation assay

HeLa cells that were either uninfected or infected with mScarlet-expressing *C. trachomatis* serovar L2 were transfected with the HA-NFAT1(4-460)-GFP plasmid. The HA-NFAT1(4-460)-GFP plasmid was a gift from Anjana Rao (Addgene plasmid # 11107 ;; RRID:Addgene_11107) [50]. At the desired time post infection, cells were washed twice with PBS, and then incubated in Ca^2+^ -free Ringer’s solution containing 1 ug/mL Hoechst stain for 30 min. Buffer was then exchanged with either DMSO or 2 μM TGCa^2+^-free Ringer’s solution. Immediately following treatment, an image was acquired using the Nikon Ti2e with a CFI60 Super Plan Fluor Phase Contrast ADM ELWD 40x Objective Lens. ND acquisition was programmed to acquire images from 4 locations per well for each condition. After incubating cells with either DMSO or TG for 5 min, the solution was exchanged with the Ca^2+^-containing Ringer’s solution, and then imaged at 18 min post Ca^2+^ addition. Imaging was performed at 37°C, 5% CO_2_, 92% relative humidity using a stage-top incubation chamber (Okolab). Following imaging, Huygens Essential software version 20.04 (Scientific Volume Imaging) was used to stabilize frames between time points and Imaris x64 software version 9.6.0 (Oxford Instruments) was used to measure the mean fluorescence intensity of NFAT-GFP in the nucleus and the cytoplasm of individual cells before and after SOCE induction. The NFAT-GFP N/C ratio was calculated as the mean nuclear NFAT-GFP fluorescence intensity / the mean cytoplasmic NFAT-GFP fluorescence intensity.

### Statistics

Statistical analysis was conducted using Prism version 9.1.1 software for Windows (GraphPad). An unpaired Student’s T test was performed for Fura-2, AM analysis, and Kruskal-Wallis test with Dunn’s posttest was performed for Fluo-4, AM relative change, GCaMP6m relative change, and NFAT-GFP N/C ratios.

## Acknowledgements

This work was supported by the Intramural Research Program of the NIAID, NIH.

We thank Rebecca Miller for excellent technical assistance, Zoe Dimond for critical review of the manuscript, and Ryan Kissinger in the Visual and Medical Arts Unit (VMA) at Rocky Mountain Laboratories for graphic artwork.

## Extended Data

**Extended Data Fig 1.**
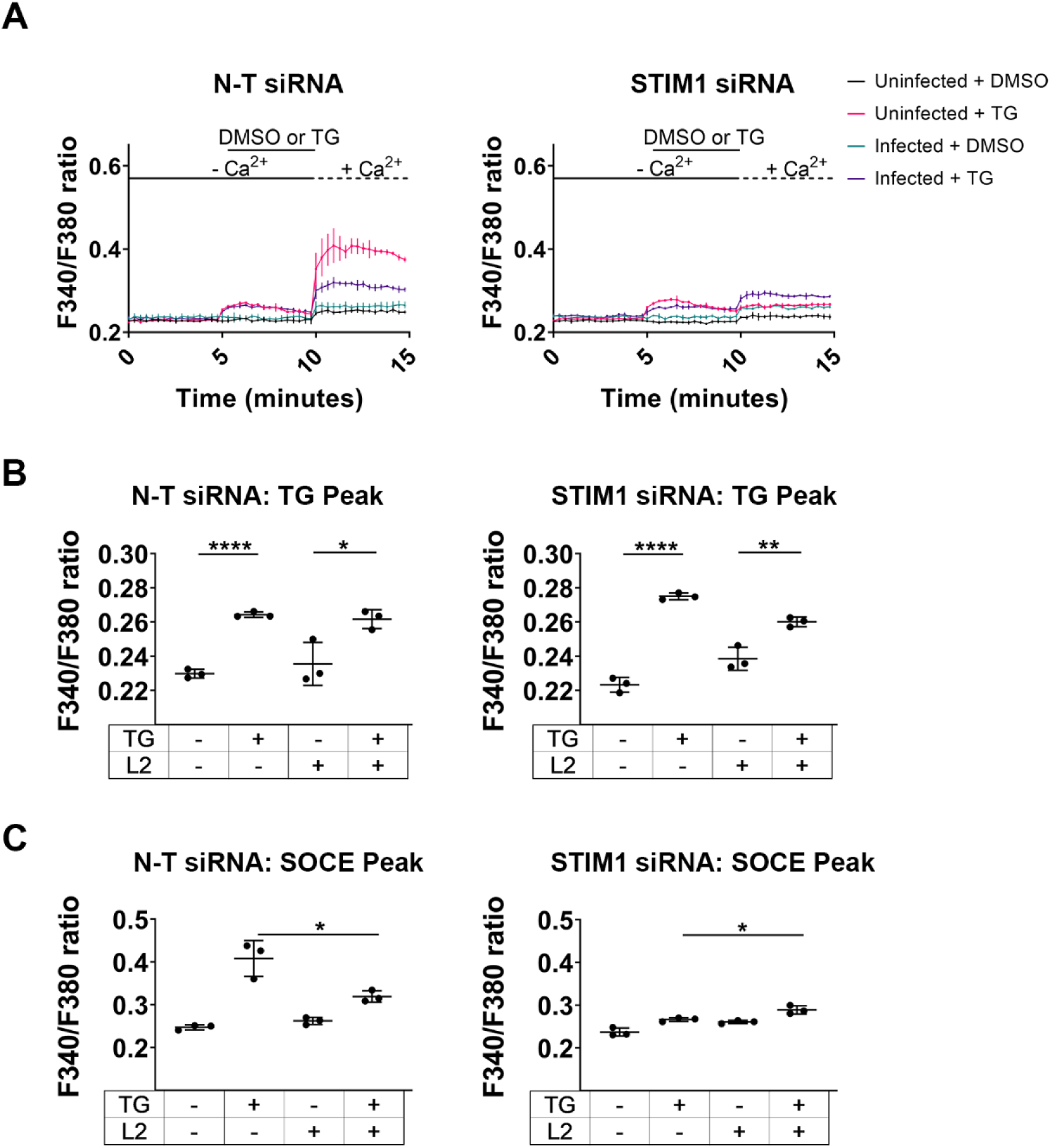
Verification that STIM1 knockdown abrogates SOCE. a A Fura-2, AM Ca^2+^ re-addition assay was performed using cells either transfected with non-targeting (N-T) siRNA or STIM1 siRNA. HeLa cells were either infected with *C. trachomatis* L2 for 24 hr or uninfected. b The relative change in [Ca^2+^]_i_ was calculated at the peak ER Ca^2+^ efflux induced by TG. Student’s T-test was used to compare the DMSO to the TG treatment, n=3. c The SOCE peak relative change in [Ca^2+^]_i_ was calculated. Student’s T-test was used to compare the uninfected TG treated condition to the 24 hpi TG treated condition, n=3. Data (a-c) are presented as mean ± SEM.

**Extended Data Fig 2.**
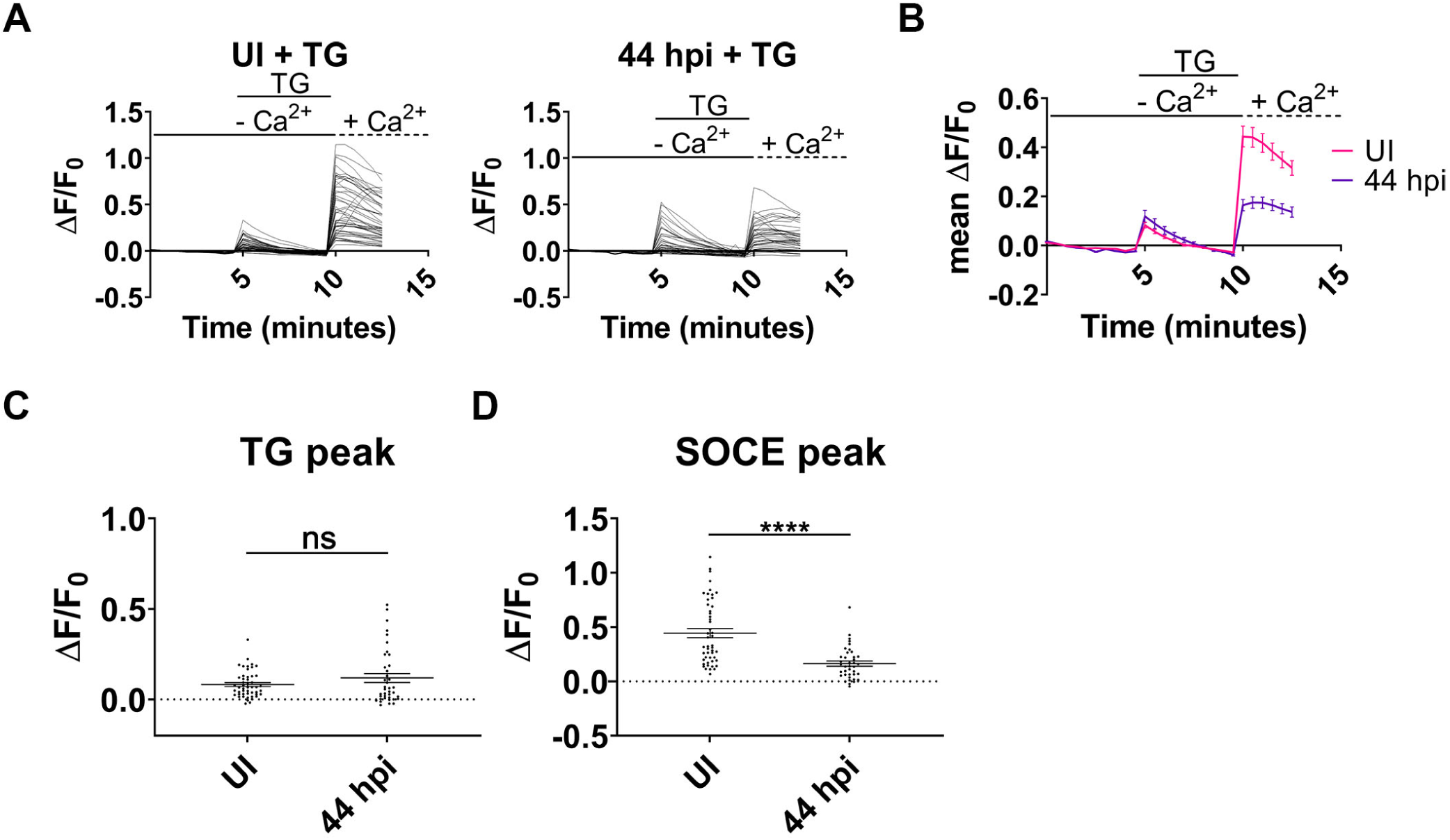
Fluo-4, AM analysis of *C. trachomatis* impairment of SOCE during late cycle development. a Single-cell analysis was performed with Fluo-4, AM-loaded HeLa cells either uninfected or infected with *C. trachomatis* L2 at 44 hpi and induced to undergo SOCE with TG. For each condition, ≥ 39 cells were analyzed. b Mean relative change in Fluo-4 fluorescence for uninfected and infected cells was calculated throughout the Ca^2+^ re-addition assay. c Single-cell analysis of the relative change in Fluo-4 fluorescence was calculated at the TG-induced peak. d Single-cell analysis of the relative change in Fluo-4 fluorescence was performed at the SOCE peak. A Student’s T-test was used for comparisons in C and D. Comparisons denoted with **** have a p value <0.0001 and ns represents no significant difference. Data (**b-d**) are presented as mean ± SEM.

**Extended Data Fig 3.**
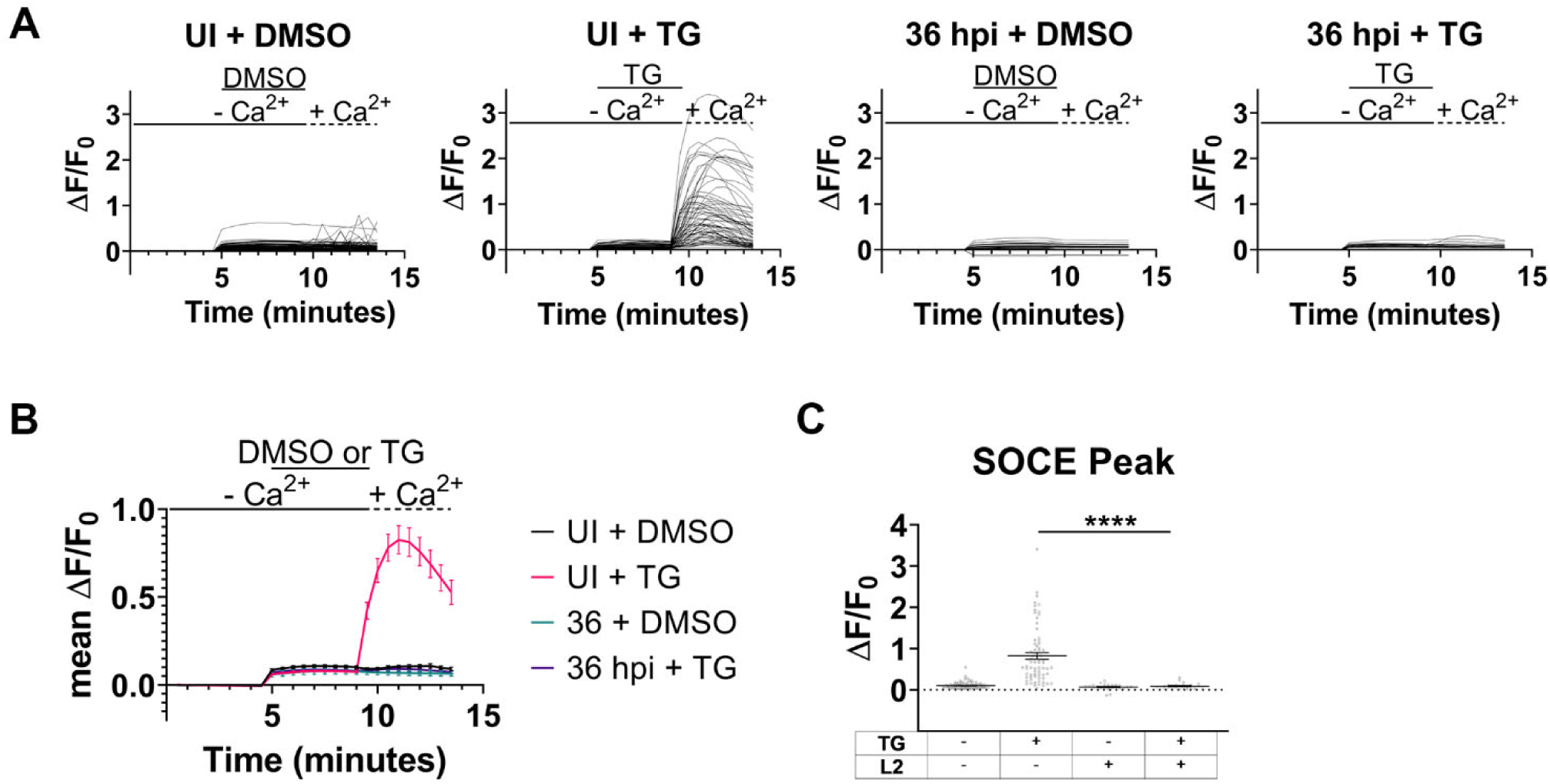
GCaMP6m analysis of *C. trachomatis* impairment of SOCE at mid-late development. a Single-cell analysis of GCaMP6m-transfected HeLa cells was conducted for either uninfected or infected with *C. trachomatis* L2 at 36 hpi. For each condition, ≥ 24 cells were analyzed. b) The mean relative change in GCaMP6m fluorescence was measured during Ca^2+^ re-addition assay. c Single-cell analysis of the relative change in GCaMP6m fluorescence was measured at the SOCE peak. A Kruskal-Wallis test was performed with Dunn’s post-hoc multiple comparisons test to compare conditions. Comparison denoted with **** has a p value <0.0001. Data (**b**,**c**) are presented as mean ± SEM.

**Extended Data Fig 4:**
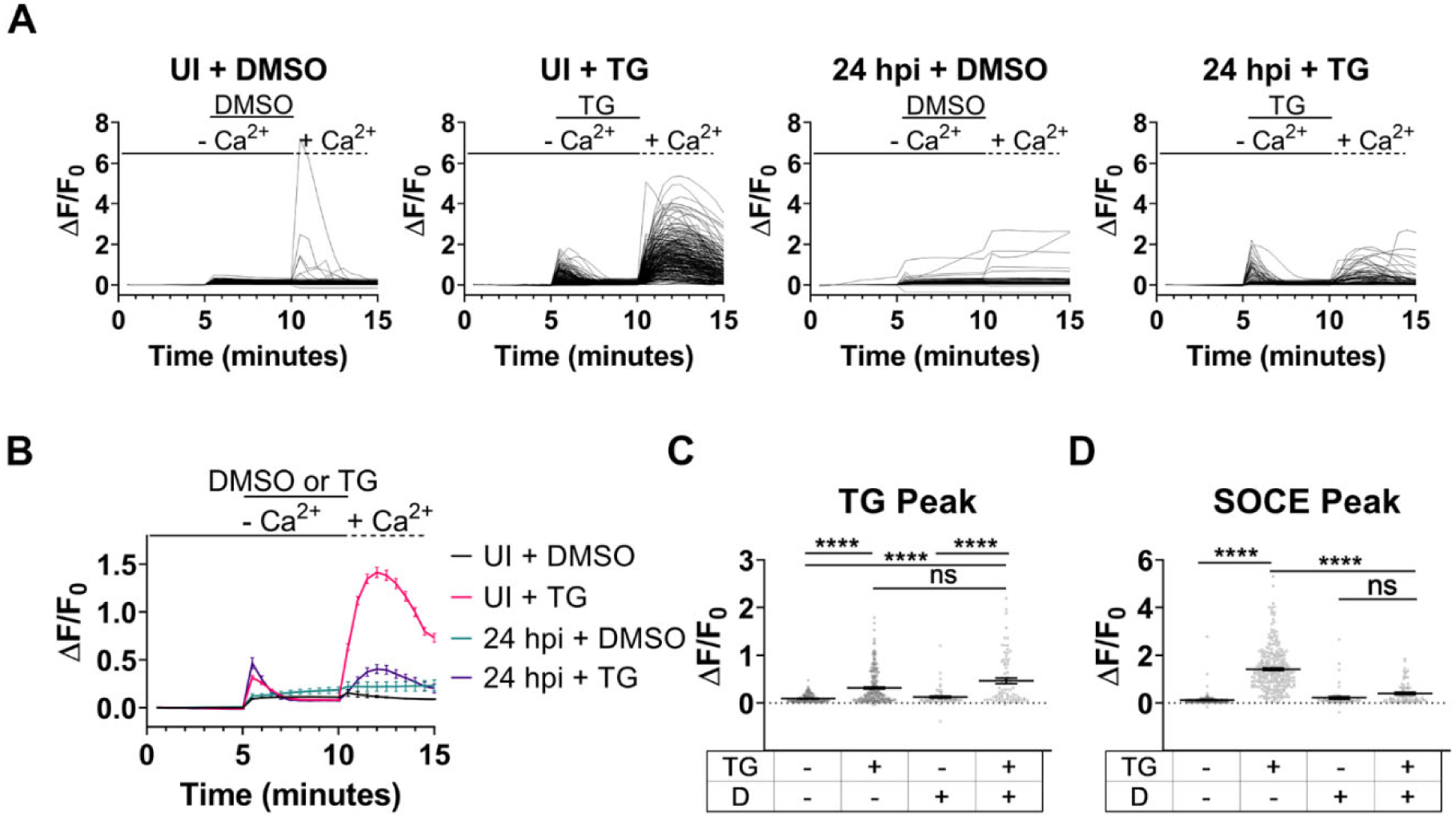
*C. trachomatis* serovar D impairs SOCE. Live-cell microscopy of GCaMP6m was used to assess [Ca^2+^]_i_ changes within HeLa cells either uninfected or infected with *C. trachomatis* serovar D. For each condition, ≥ 69 cells were analyzed. a Single-cell analysis of relative change in GCaMP6m fluorescence was conducted throughout the Ca^2+^ re-addition assay. b The mean relative change in GCaMP6m fluorescence was measured for cells from (**a**). c The relative change in GCaMP6m fluorescence at the TG peak was calculated for each condition. d The relative change in GCaMP6m fluorescence was assessed for the SOCE peak. Analyzes were performed using Kruskal-Wallis test with Dunn’s post-hoc multiple comparisons test (**c**,**d**). Comparisons denoted with **** have a p value <0.0001 and ns represents no significant difference. Data (**b-d**) are presented as mean ± SEM.

## Supplementary Information

**Supplementary Video 1. Time series of NFAT-GFP nuclear localization in uninfected cells**

HeLa cells transfected with NFAT-GFP were treated with 2 μM TG in Ca^2+^-free Ringer’s solution then solution was exchanged with Ca^2+^-containing Ringer’s solution to induce SOCE. Time series was acquired by imaging cells every minute for 5 minutes in the Ca^2+^-free Ringer’s solution, and then every minute for 20 minutes in Ca^2+^-containing Ringer’s solution.

**Supplementary Video 2. Time series of NFAT-GFP nuclear localization in *C. trachomatis* infected cells**

HeLa cells were transfected with NFAT-GFP and infected with mScarlet-expressing *C. trachomatis* L2 for 24 hours. Cell treatments and imaging was performed as described in Supplementary Video 1.

## References

1. Abdelrahman, Y. M. and Belland, R. J. The chlamydial developmental cycle. FEMS Microbiol Rev 29: 949–959. (2005).

2. Moulder, J. W. Interaction of chlamydiae and host cells in vitro. Microbiol Rev 55: 143–190. (1991).

3. Wyrick, P. B. Intracellular survival by Chlamydia. Cell Microbiol 2: 275–282. (2000).

4. Schachter, J. Infection and disease epidemiology. In Chlamydia; Intracellular biology, pathogenesis, and immunity, R.S. Stephens, (ed.), ASM Press: Washington, D.C. pp. 139–169. 1999.

5. Burton, M. J. Trachoma: an overview. Br. Med. Bull. 84: 99–116. (2007).

6. Abdelsamed, H., Peters, J., and Byrne, G. I. Genetic variation in Chlamydia trachomatis and their hosts: impact on disease severity and tissue tropism. Future Microbiol 8: 1129–1146. (2013).

7. Betts-Hampikian, H. J. and Fields, K. A. The chlamydial type III secretion mechanism: revealing cracks in a tough nut. Front. Microbiol. 1: 114. (2010).

8. Bannantine, J. P., Griffiths, R. S., Viratyosin, W., Brown, W. J., and Rockey, D. D. A secondary structure motif predictive of protein localization to the chlamydial inclusion membrane. Cell Microbiol 2: 35–47. (2000).

9. Weber, M. M., Bauler, L. D., Lam, J., and Hackstadt, T. Expression and localization of predicted inclusion membrane proteins in Chlamydia trachomatis. Infect Immun (2015).

10. Rockey, D. D., Grosenbach, D., Hruby, D. E., Peacock, M. G., Heinzen, R. A., and Hackstadt, T. Chlamydia psittaci IncA is phosphorylated by the host cell and is exposed on the cytoplasmic face of the developing inclusion. Mol Microbiol 24: 217–228. (1997).

11. Hackstadt, T., Scidmore-Carlson, M. A., Shaw, E. I., and Fischer, E. R. The Chlamydia trachomatis IncA protein is required for homotypic vesicle fusion. Cell Microbiol 1: 119–130. (1999).

12. Scidmore, M. A. and Hackstadt, T. Mammalian 14-3-3beta associates with the Chlamydia trachomatis inclusion membrane via its interaction with IncG. Mol Microbiol 39: 1638–1650. (2001).

13. Mital, J., Miller, N. J., Fischer, E. R., and Hackstadt, T. Specific chlamydial inclusion membrane proteins associate with active Src family kinases in microdomains that interact with the host microtubule network. Cell Microbiol 12: 1235–1249. (2010).

14. Lutter, E. I., Barger, A. C., Nair, V., and Hackstadt, T. Chlamydia trachomatis inclusion membrane protein CT228 recruits elements of the myosin phosphatase pathway to regulate release mechanisms. Cell Rep 3: 1921–1931. (2013).

15. Dimond, Z. E., Suchland, R. J., Baid, S., LaBrie, S. D., Soules, K. R., Stanley, J., Carrell, S., Kwong, F., Wang, Y., Rockey, D. D., Hybiske, K., and Hefty, P. S. Inter-species lateral gene transfer focused on the Chlamydia plasticity zone identifies loci associated with immediate cytotoxicity and inclusion stability. Mol Microbiol 116: 1433–1448. (2021).

16. Mital, J., Lutter, E. I., Barger, A. C., Dooley, C. A., and Hackstadt, T. Chlamydia trachomatis inclusion membrane protein CT850 interacts with the dynein light chain DYNLT1 (Tctex1). Biochem Biophys Res Commun 462: 165–170. (2015).

17. Nguyen, P. H., Lutter, E. I., and Hackstadt, T. Chlamydia trachomatis inclusion membrane protein MrcA interacts with the inositol 1,4,5-trisphosphate receptor type 3 (ITPR3) to regulate extrusion formation. PLoS Pathog 14: e1006911. (2018).

18. Berridge, M. J., Lipp, P., and Bootman, M. D. The versatility and universality of calcium signalling. Nat Rev Mol Cell Biol 1: 11–21. (2000).

19. Graef, I. A., Chen, F., and Crabtree, G. R. NFAT signaling in vertebrate development. Curr Opin Genet Dev 11: 505–512. (2001).

20. Berridge, M. J., Bootman, M. D., and Roderick, H. L. Calcium signalling: dynamics, homeostasis and remodelling. Nat. Rev. Mol. Cell Biol. 4: 517–529. (2003).

21. Prakriya, M. and Lewis, R. S. Store-Operated Calcium Channels. Physiol Rev 95: 1383–1436. (2015).

22. Venkatachalam, K., van Rossum, D. B., Patterson, R. L., Ma, H. T., and Gill, D. L. The cellular and molecular basis of store-operated calcium entry. Nat. Cell Biol. 4: E262–272. (2002).

23. Agaisse, H. and Derré, I. Expression of the effector protein IncD in Chlamydia trachomatis mediates recruitment of the lipid transfer protein CERT and the endoplasmic reticulum-resident protein VAPB to the inclusion membrane. Infect. Immun. 82: 2037–2047. (2014).

24. Stanhope, R., Flora, E., Bayne, C., and Derré, I. IncV, a FFAT motif-containing Chlamydia protein, tethers the endoplasmic reticulum to the pathogen-containing vacuole. Proc. Natl. Acad. Sci. U.S.A. 114: 12039–12044. (2017).

25. Agaisse, H. and Derré, I. STIM1 Is a novel component of ER-Chlamydia trachomatis inclusion membrane contact sites. PLoS ONE 10: e0125671. (2015).

26. Bird, G. S., DeHaven, W. I., Smyth, J. T., and Putney, J. W. J. Methods for studying store-operated calcium entry. Methods 46: 204–212. (2008).

27. Flourakis, M., Van Coppenolle, F., Leheńkyi, V., Beck, B., Skryma, R., and Prevarskaya, N. Passive calcium leak via translocon is a first step for iPLA2-pathway regulated store operated channels activation. FASEB J. 20: 1215–1217. (2006).

28. Van Coppenolle, F., Vanden Abeele, F., Slomianny, C., Flourakis, M.,., Hesketh, J., Dewailly, E., and Prevarskaya, N. Ribosome-translocon complex mediates calcium leakage from endoplasmic reticulum stores.. J. Cell Sci. 117: 135–142. (2004).

29. Nakai, J., Ohkura, M., and Imoto, K. A high signal-to-noise Ca(2+) probe composed of a single green fluorescent protein. Nat. Biotechnol. 19: 137–141. (2001).

30. Clapham, D. E. Calcium signaling. Cell 131: 1047–1058. (2007).

31. Rao, A., Luo, C., and Hogan, P. G. Transcription factors of the NFAT family: regulation and function. Annu. Rev. Immunol. 15: 707–747. (1997).

32. Bootman, M. D., Thomas, D., Tovey, S. C., Berridge, M. J., and Lipp, P. Nuclear calcium signalling. Cell Mol Life Sci 57: 371–378. (2000).

33. Crabtree, G. R. and Olson, E. N. NFAT signaling: choreographing the social lives of cells. Cell 109 Suppl: S67–79. (2002).

34. Hybiske, K. and Stephens, R. S. Mechanisms of host cell exit by the intracellular bacterium Chlamydia. Proc Natl Acad Sci USA 104: 11430–11435. (2007).

35. Derre, I., Swiss, R., and Agaisse, H. The lipid transfer protein CERT interacts with the Chlamydia inclusion protein IncD and participates to ER-Chlamydia inclusion membrane contact sites. PLoS Pathog 7: e1002092. (2011).

36. Dumoux, M., Clare, D. K., Saibil, H. R., and Hayward, R. D. Chlamydiae assemble a pathogen synapse to hijack the host endoplasmic reticulum. Traffic 13: 1612–1627. (2012).

37. Béliveau, É., Lessard, V., and Guillemette, G. STIM1 positively regulates the Ca2+ release activity of the inositol 1,4,5-trisphosphate receptor in bovine aortic endothelial cells. PLoS ONE 9: e114718. (2014).

38. Quayle, A. J. The innate and early immune response to pathogen challenge in the female genital tract and the pivotal role of epithelial cells. J Reprod Immunol 57: 61–79. (2002).

39. Buckner, L. R., Lewis, M. E., Greene, S. J., Foster, T. P., and Quayle, A. J. Chlamydia trachomatis infection results in a modest pro-inflammatory cytokine response and a decrease in T cell chemokine secretion in human polarized endocervical epithelial cells. Cytokine 63: 151–165. (2013).

40. Jairaman, A., Yamashita, M., Schleimer, R. P., and Prakriya, M. Store-operated Ca2+ release-activated Ca2+ channels regulate PAR2-activated Ca2+ signaling and cytokine production in airway epithelial cells. J Immunol 195: 2122–2133. (2015).

41. Pan, H. Y., Ladd, A. V., Biswal, M. R., and Valapala, M. Role of nuclear factor of activated T cells (NFAT) pathway in regulating autophagy and inflammation in retinal pigment epithelial cells. Int J Mol Sci 22: (2021).

42. Yokoyama, K., Higashi, H., Ishikawa, S., Fujii, Y., Kondo, S., Kato, H., Azuma, T., Wada, A., Hirayama, T., Aburatani, H., and Hatakeyama, M. Functional antagonism between Helicobacter pylori CagA and vacuolating toxin VacA in control of the NFAT signaling pathway in gastric epithelial cells. Proc Natl Acad Sci U S A 102: 9661–9666. (2005).

43. Blatter, L. A. Tissue specificity: SOCE: implications for Ca(2+) handling in endothelial cells. Adv Exp Med Biol 993: 343–361. (2017).

44. Miskin, J. E., Abrams, C. C., Goatley, L. C., and Dixon, L. K. A viral mechanism for inhibition of the cellular phosphatase calcineurin. Science 281: 562–565. (1998).

45. Zhou, Y., Heesom, K., Osborn, K., AlMohammed, R., Sweet, S. M., and Sinclair, A. J. Identifying the cellular interactome of Epstein-Barr Virus lytic regulator Zta reveals cellular targets contributing to viral replication. J Virol 94: (2020).

46. Szabò, I., Brutsche, S., Tombola, F., Moschioni, M., Satin, B., Telford, J. L., Rappuoli, R., Montecucco, C., Papini, E., and Zoratti, M. Formation of anion-selective channels in the cell plasma membrane by the toxin VacA of Helicobacter pylori is required for its biological activity. EMBO J 18: 5517–5527. (1999).

47. Caldwell, H. D., Kromhout, J., and Schachter, J. Purification and partial characterization of the major outer membrane protein of Chlamydia trachomatis. Infect Immun 31: 1161–1176. (1981).

48. Furness, G., Graham, D. M., and Reeve, P. The titration of trachoma and inclusion blennorrhoea viruses in cell cultures. J. Gen. Microbiol. 23: 613–619. (1960).

49. Chen, T. W., Wardill, T. J., Sun, Y., Pulver, S. R., Renninger, S. L., Baohan, A., Schreiter, E. R., Kerr, R. A., Orger, M. B., Jayaraman, V., Looger, L. L., Svoboda, K., and Kim, D. S. Ultrasensitive fluorescent proteins for imaging neuronal activity. Nature 499: 295–300. (2013).

50. Aramburu, J., Yaffe, M. B., López-Rodríguez, C., Cantley, L. C., Hogan, P. G., and Rao, A. Affinity-driven peptide selection of an NFAT inhibitor more selective than cyclosporin A. Science 285: 2129–2133. (1999).

